# Measuring the relationship between sleep, physical activity and cognition

**DOI:** 10.1101/599092

**Authors:** Marta Swirski, Netasha Shaikh, Amy Chinner, Ellen Gaaikema, Elizabeth Coulthard

## Abstract

Biochemical and neuropsychological changes due to poor sleep may contribute to the development of neurodegenerative disorders, such as dementia. Physical activity is widely thought to improve sleep; however, the optimal intensity/duration of physical activity required is unknown. This 14-week, single-blind study (n=23) investigated the feasibility of a self-directed physical activity intervention in healthy adults using actigraphy and cognitive function measures as primary outcomes. Participants were randomised to a control group (no change in routine) or the intervention group (increased physical activity) and were provided with an actigraphy device to monitor activity. Participants completed daily sleep/activity diaries and three cognitive assessment sessions. Vigorous physical activity increased between baseline and week 3 for the intervention group only, with no identifiable impact on sleep. This change was not sustained at week 12. Performance on an executive function task and delayed visuospatial recall improved from baseline to week 12 for the intervention group only. Contrary to our expectations, increasing light-moderate physical activity was associated with more impaired sleep across all participants. It is clear that the relationships between physical activity, sleep and cognition are complex and require further investigation. We discuss optimal methodologies for clinical trials investigating physical activity and/or sleep interventions targeting cognition.

## Introduction

Dementia is a highly prevalent, and currently irreversible, syndrome affecting approximately 35 million people worldwide^1^. Increasing evidence suggests that deficient or dysregulated sleep, including insomnia, sleep disordered breathing, excessive daytime sleepiness and circadian rhythm sleep disorder, is commonly associated with neurodegenerative diseases^2^. Over 60% of patients diagnosed with mild cognitive impairment or Alzheimer’s disease (AD) report sleep disturbance^2,3^. Putatively, this sleep disturbance is both caused, and exacerbated, by dementia-related pathologies in the brain^4,5^.

The biochemical mechanisms through which sleep disturbances may contribute to dementia are still to be established. In a murine model of AD, chronic sleep deprivation exacerbated memory impairment, senile plaque deposition and phosphorylated tau levels^6^. In a separate study, sleep-deprived rats had long-lasting neurochemical (noradrenaline and dopamine) changes^7^. This potential role of disturbed sleep in the aetiology of dementia raises the possibility of sleep as a modifiable risk factor to target disease processes in the brain. Furthermore, a large number of studies have provided evidence supporting the role of sleep in memory processing and brain plasticity^8,9,10,11,12,13,14^ Therefore, in addition to improved sleep potentially slowing the progression of dementia, a direct cognitive enhancement may also be possible.

Exercise has long been associated with better sleep outcomes and is often recommended as a nonpharmacological treatment option for people experiencing sleep disorders^15,16,17^. In addition, regular exercise focusing on functional fitness, such as walking, has been associated with significant reductions in dependence and disability in older adults ^18,19^ as well as benefits on cognition^20,21,22^. However, there are still no established guidelines providing a structured approach for implementing these lifestyle changes, especially for people experiencing cognitive impairment or dementia. Several reasons for this include inconsistent evidence regarding the benefit of exercise in this patient population as well as difficulties in drawing conclusions from studies using different study designs and methodologies.

The complex process of translating a person’s sleep and physical activity into quantifiable and comparable measures remains a challenge. Consequently, the importance of applying appropriate methodology is of critical importance when capturing these data. Both subjective and objective methods have been explored, including self-reported questionnaires (Pittsburgh Sleep Quality Index (PSQI)^23^; Consensus Sleep Diaries^24^) and, more recently, polysomnography and actigraphy. Studies investigating changes in people with diagnosed dementia tend to focus on broad cognitive outcomes such as performance on standardised assessments, for example, the Repeatable Battery for the Assessment of Neuropsychological Status (RBANS) and Alzheimer’s Disease Assessment Scale-cognitive subscale (ADAScog), or markers of functional cognitive state, such as the Clinical Dementia Rating (CDR). As these measures do not focus on the specific cognitive effects of sleep and exercise, changes may only become apparent after many months, or years, of follow-up. Therefore, administration of measure that can probe both sleep and physical activity, like actigraphy, may provide a more useful initial outcome measure to assess the effectiveness of interventions to improve sleep and delay dementia. Although the use of actigraphy devices in measuring sleep and/or physical activity is expanding in research, there is currently limited information about the functionality of this technology, particularly for measuring activity in older and cognitively impaired populations.

We hypothesise that 1) increased physical activity enhances sleep quality 2) an intervention that increases levels of physical activity will improve sleep and 3) improved sleep quality enhances cognition. In this study, we aim to explore the best way to test these hypotheses by initially conducting a feasibility study using an actigraphy device to measure sleep and physical activity in healthy adults. In addition, we aim to evaluate the relationships between physical activity, sleep and cognition with a view to developing more sensitive outcome measures for future interventional sleep trials.

## Results

### Recruitment

We screened 144 volunteers and 17 volunteers from the Join Dementia Research website and the ReMemBr group healthy volunteer database, respectively (total screened, n=161). The total number of volunteers consenting to take part in this study was 58, of which 19 withdrew (most withdrawals took place following distribution of the actigraphy monitor at week 6 for reasons unspecified and 2 withdrawals were due to injury unrelated to study participation), 7 were lost to follow-up and 5 were ineligible. A total of 27 participants completed the study of which 23 were evaluable for analysis (n=12, exercise group; n=11, control group). Four participants were ineligible for analysis due to insufficient wearing of the actigraphy monitor/missing data (**Figure 1**). Participant demographics for those included in the analysis are presented in **Table 1**.

**Figure 1.**
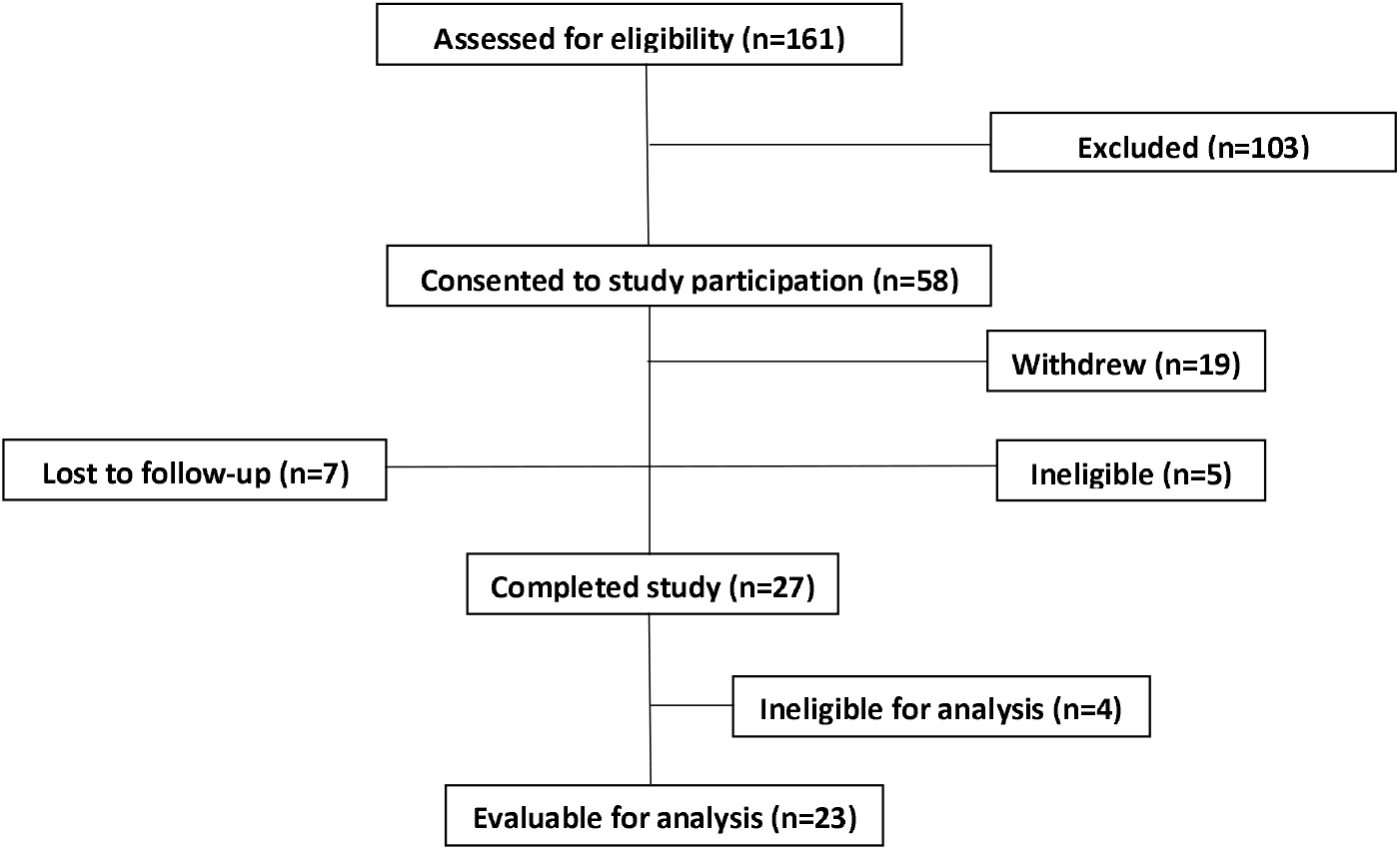
We screened a total number of 161 volunteers. The total number of volunteers consenting to take part in this study was 58 of which 19 withdrew, 7 were lost to follow-up and 5 were ineligible. A total of 27 participants completed the study of which 23 were evaluable for analysis. Four participants were ineligible for analysis.

**Table 1.** Demographics of participants enrolled into the control (n=11) and intervention (n=12) groups. No significant differences were found between groups with respect to age, height, weight, years of education, gender, Depression/Anxiety/Stress Scale (DASS) score, Geriatric depression scale (GDS) score, systolic/diastolic blood pressure, heart rate, Montreal Cognitive Assessment (MoCA) score or peak flow reading at baseline (first participant visit).

|  | Control (n=11) |  | Intervention (n=12) |  |  |
| --- | --- | --- | --- | --- | --- |
|  | Mean | Standard deviation (SD) | Mean | Standard deviation (SD) | Significant (S) vs non-significant (NS) |
| Age | 60.2 | 13.7 | 52.8 | 13.8 | NS |
| Height (cm) | 168.1 | 7.0 | 168.5 | 12.2 | NS |
| Weight (kg) | 70.0 | 7.0 | 69.6 | 16.0 | NS |
| Years of education (years) | 15.9 | 3.9 | 17.3 | 2.7 | NS |
| % Female | 45.45 | - | 58.3 | - | NS |
| DASS score (depression) | 1.0 | 1.6 | 1.4 | 2.7 | NS |
| DASS score (anxiety) | 0.6 | 0.9 | 1.7 | 2.3 | NS |
| DASS score (stress) | 4.9 | 5.1 | 5.9 | 3.5 | NS |
| Geriatric Depression Scale score | 1.1 | 1.1 | 1.7 | 2.9 | NS |
| Systolic blood pressure (mmHg) | 132.0 | 10.9 | 130.5 | 17.1 | NS |
| Diastolic blood pressure (mmHg) | 77.8 | 12.1 | 80.9 | 12.2 | NS |
| Heart rate (bpm) | 64.4 | 11.9 | 71.1 | 10.1 | NS |
| MoCA score | 27.9 | 1.9 | 27.9 | 1.6 | NS |
| Peak flow reading (l/min) | 495.1 | 98.7 | 425.4 | 98.7 | NS |

### Feasibility and acceptability

As mentioned above, 4 participants who completed the study had significantly high non-wear time (>50%) so were excluded from the main analysis. Participants in both the intervention and control groups completed sleep and exercise diaries for the entire study (with some blank entries due to forgetfulness). Several participants reported that completion of diaries became monotonous. No serious adverse events occurred during the study.

### Physical Activity: modest, unsustained effect of intervention on physical activity compared with baseline

#### % of time spent sedentary

Repeated measures ANOVA, with epoch as within subject variable (baseline vs week 1 & 2 and baseline vs week 11 & 12), intervention group as between subject factor and age as a covariate, demonstrated no significant interaction between participant group and study epoch when measuring % of time spent sedentary for baseline vs week 1 & 2 (F (1, 19) = 2.132, p = 0.161) or baseline vs week 11 & 12 (F (1,19) = 1.412, p = 0.251) **(Figure 2A) (Table 2 for respective effect size data)**.

**Figure 2.**
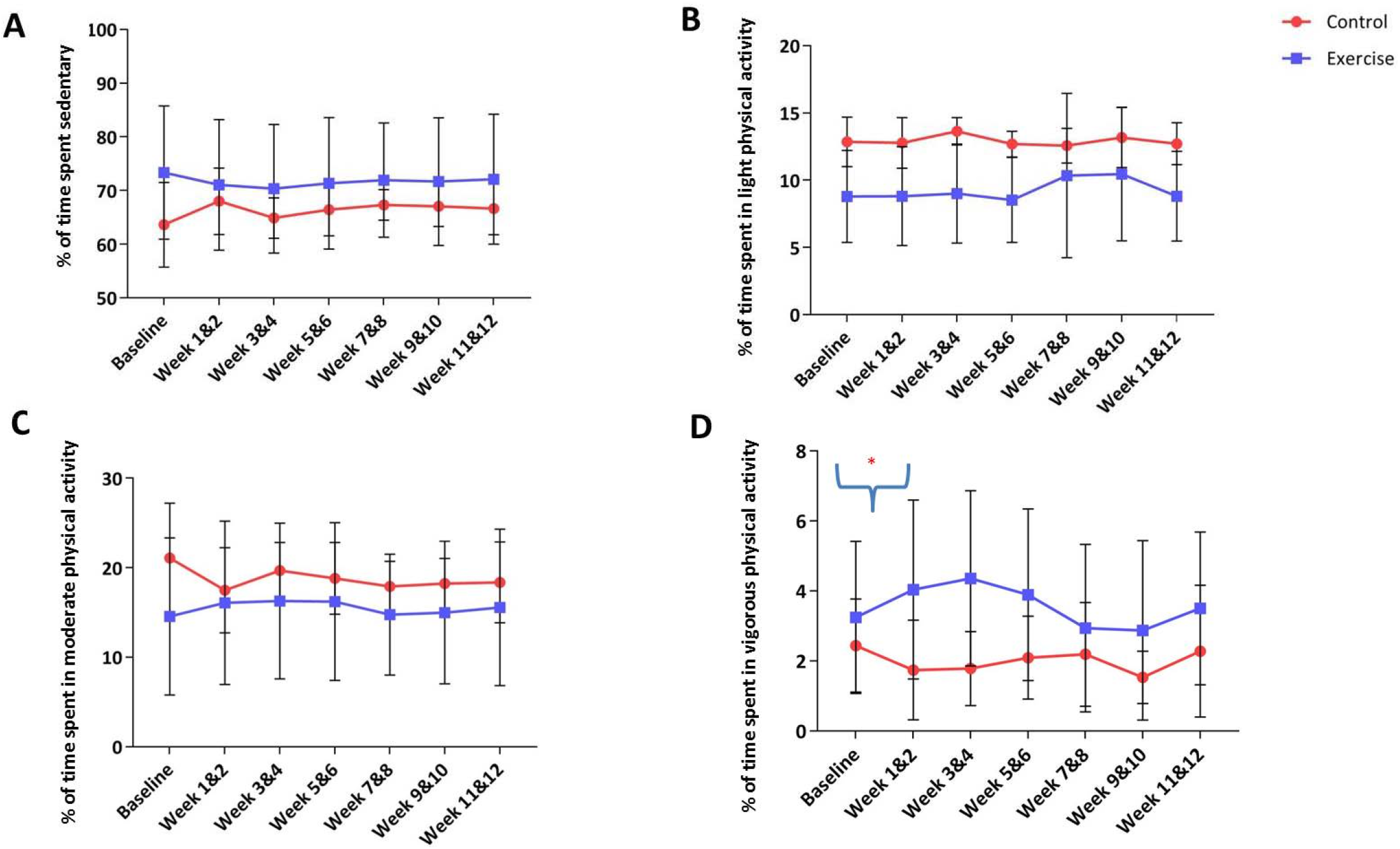
Mean % of time spent sedentary (A), mean % of time spent in light physical activity (B) mean % of time spent in moderate physical activity (C) and mean % of time in vigorous physical activity (D) in control (red) and exercise (blue) groups over 14 weeks. Error bars indicate SD. A) No interaction between participant group was observed with mean % spent sedentary at week 1 & 2 and week 11 & 12 vs baseline (F (1,19) = 2.132, p = 0.161; (F (1,19) = 1.412, p = 0.251, respectively) B) No interaction between participant group was observed with mean % of time light physical activity at week 1 & 2 and week 11 & 12 vs baseline (F (1, 19) = 0.526, p = 0.478; (F (1, 19) = 0.686, p = 0.420, respectively) C) No interaction between participant group was observed with mean % of time in moderate physical activity at week 1 & 2 and week 11 & 12 vs baseline (F (1,19) = 1.815, p = 0.195; F (1,19) = 0.487, p = 0.495, respectively) D) A significant interaction between participant group and study epoch measuring % of time spent in vigorous physical activity for baseline vs week 1 & 2 (F (1,19) = 9.512, p = 0.006*) but no significant results for baseline vs week 11 & 12 (F (1,19) = 0.037, p = 0.849) was observed.

**Table 2.** Partial ETA square values with epoch as within subject variable (baseline vs week 1&2 and baseline vs week 11&12), intervention group as between subject factor and age as a covariate. Significant findings (determined by repeated measure ANOVA) highlighted in red.

|  | % sedentary | % light exercise | % moderate exercise | % vigorous exercise |
| --- | --- | --- | --- | --- |
| Baseline vs week 1 & 2 | .101 | .028 | .092 | .346 |
| Baseline vs week 11 & 12 | .077 | .041 | .030 | .002 |

#### % of time spent in light physical activity

Repeated measure ANOVA, with epoch as within subject variable (baseline vs week 1 & 2 and baseline vs week 11 & 12), intervention group as between subject factor and age as a covariate, demonstrated no significant interaction between participant group and study epoch when measuring % of time spent in light physical activity for baseline vs week 1 & 2 (F (1,19) = 0.526, p = 0.478) or baseline vs week 11 & 12 (F (1, 19) = 0.686, p = 0.420) **(Figure 2B) (Table 2 for respective effect size data)**.

#### % of time spent in moderate physical activity

Repeated measure ANOVA, with epoch as within subject variable (baseline vs week 1 & 2 and baseline vs week 11 & 12), intervention group as between subject factor and age as a covariate, demonstrated no significant interaction between participant group and study epoch when measuring % of time spent in moderate exercise for baseline vs week 1 & 2 (F (1, 19) = 1.815, p = 0.195) or baseline vs week 11 & 12 (F (1, 19) = 0.487, p = 0.495) **(Figure 2C) (Table 2 for respective effect size data)**.

#### % of time spent in vigorous physical activity

Repeated measure ANOVA, with epoch as within subject variable (baseline vs week 1 & 2 and baseline vs week 11 & 12), intervention group as between subject factor and age as a covariate, demonstrated a significant interaction between participant group and study epoch when measuring % of time spent in vigorous physical activity for baseline vs week 1 & 2 (F (1, 19) = 9.512, p = 0.006) but no significant results for baseline vs week 11 & 12 (F (1,19) = 0.037, p = 0.849) **(Figure 2D) (Table 2 for respective effect size data)**.

### Sleep: no effect of intervention on sleep compared with baseline

Repeated measure ANOVA, with epoch as within subject variable (baseline vs week 1 & 2 and baseline vs week 11 & 12), intervention group as between subject factor and age as a covariate, demonstrated no significant interaction between participant group and study epoch when measuring total sleep time, sleep efficiency, or number of awakenings for baseline vs week 1 & 2 or baseline vs week 11 & 12 (Total sleep time: (F (1, 20) = 0.226, p = 0.640) for baseline vs week 1 & 2 and (F (1,16) = 1.326, p = 0.266) for baseline vs week 11 & 12; Sleep efficiency: (F (1, 20) = 1.327, p = 0.263) for baseline vs week 1 & 2 and (F (1,16) = 1.783, p = 0.200) for baseline vs week 11 & 12; Number of awakenings: (F (1, 20) = 0.171, p = 0.683) for baseline vs week 1 & 2 and (F (1,16) = 0.328, p = 0.575)) for baseline vs week 11 & 12.

### Cognition: significant effect of physical activity intervention on Tower of London (ToL) and Brief Visuospatial Memory Test-revised (BVMT-R) performance only

Repeated measure ANOVA with age as covariate and participant group as between subjects variable were performed to look for the effect of intervention on cognitive performance. Results demonstrated no effect of intervention (baseline week 0 vs end week 12) on MoCA score (F (1, 20) = 2.034, p = 0.169), Maniken test (F (1,18) = 3.4, p = 0.085), Reaction speed (F 1, 16) = 0.147, p = 0.706) or HVLT (F (1, 18) = 0.46, p = 0.833). A significant effect of intervention on ToL (F (1, 18) = 6.104, p = 0.024, improvement in performance) and BVMT-R delayed recall score (F (1, 20) = 4.338, p = 0.050, improvement in performance) was observed. Respective partial ETA values are presented in **Table 3**.

**Table 3.** Partial ETA square values with epoch as within subject variable (week 0 (baseline) vs week 12 (end of study)), intervention

|  | MoCA | Maniken | Tower of London | Reaction speed | HVLT (trial 4) | BVMT (delayed recall) |
| --- | --- | --- | --- | --- | --- | --- |
| Week 0 vs week 12 | .092 | .185 | .253 | .009 | .002 | .178 |

Next, we sought to explore the relationships between physical activity, sleep and cognitive outcomes. As the effects of the intervention were modest we decided to pool the intervention and control groups together for further analysis.

### Physical activity and sleep: significant correlations between physical activity and indicators of sleep disruption

Linear regression was performed to seek predictors of sleep efficiency, number of awakenings and total sleep time at baseline, week 1 & 2 and week 11 & 12. All results were corrected for age and multiple comparisons. At baseline, weeks 1 & 2 and weeks 11 & 12, a significant positive relationship between % of light physical activity conducted and number of awakenings was found (baseline: Pearson r = 0.575, p = 0.01; weeks 1 & 2: Pearson r = 0.575, p = 0.01; weeks 11 & 12: Pearson r = 0.546, p = 0.0023). No other significant findings were found between physical activity and sleep at baseline. At weeks 1 & 2, a significant negative relationship between % of moderate physical activity conducted and sleep efficiency (Pearson r = −0.707, p = 0.001), as well as a significant positive relationship between % of moderate physical activity conducted and number of awakenings (Pearson r = 0.525, p = 0.021), was found. At study end (weeks 11 & 12), a significant positive relationship was found between % of time spent sedentary and total sleep time (Pearson r = 0.495, p = 0.044). In addition, a significant negative relationship was found between % of moderate physical activity conducted and sleep efficiency (Pearson r = −0.639, p = 0.006), and a significant positive relationship with number of awakenings (Pearson r = 0.545, p = 0.024).

### Cognitive outcomes: MoCA score predicted by a combination of age, total sleep time, sleep efficiency and moderate physical activity

Linear regressions were performed to see if any representative physical activity (moderate or vigorous activity), sleep measures (total sleep time and sleep efficiency) or age predicted cognition at baseline. These measures were chosen as representative rather than using all possible measures to avoid the confound of multi-collinearity, therefore, measures were chosen with as little theoretical overlap as possible. MoCA score significantly predicted by a combination of age, total sleep time, sleep efficiency and moderate physical activity (F(4,16)=3.32, p=0.036) with no individual variable being a significant predictor.

### Sleep measures: relationship between actigraph-measured sleep efficiency and self-reported quality of sleep

As we were interested in establishing the best methodologies for the investigation of sleep, physical activity and cognition, we conducted a correlation analysis between self-reported quality of sleep (scale 1-5), assessed by participants through a sleep diary, and sleep efficiency, assessed by the actigraph. There was a significant positive relationship between self-reported sleep quality and actigraph-measured sleep efficiency (Spearman r = 0.2031, p < 0.0001).

## Discussion

One aim of our investigation was to explore the feasibility of a simple, self-directed, intervention to increase physical activity in people across a spectrum of ages, including those at increased risk of dementia. Our first finding was that the intervention led to an increase in vigorous physical activity in the first two weeks, but that this change was not sustained at 12 weeks. It was also found that the intervention had no effect on sleep but higher average levels of light and moderate physical activity, when pooled across both groups, predicted poorer sleep as assessed by number of awakenings and to a lesser extent with sleep efficiency and total sleep time. From a cognitive perspective, the intervention was associated with improvements on the ToL and BVMT-R delayed recall cognitive tasks only. This provides some insight into the benefit of physical activity on cognitive function suggesting it may be specific to certain cognitive processes. Overall, we did not find that physical activity or sleep alone significantly predicted cognitive performance, although MoCA score was explained by a combination of age, total sleep time, sleep efficiency and moderate physical activity illustrating the complexity of these interactions.

Our first finding is that this intervention has a very weak and unsustained effect on physical activity. Reasons for this could include lack of motivation or “buy-in” of participants. We met with participants three times over the 14 weeks for physical/cognitive assessments in which we enquired generally about their progress; however, the study team were blinded to study group and so detailed feedback could not be provided. Future studies could utilise more structured interaction with the participant through a questionnaire or feedback sensor technology, and triggered feedback sessions. In addition, future studies could tailor the intervention regimen based on baseline assessment of both physical activity and Patient Activation Measures^25^. We would also utilise devices with better battery life, or home charging capabilities, with monitoring over a shorter time period to limit participant burden. Several participants informed us that the sleep/activity diaries were fairly tedious to complete every day for 14 weeks and one participant felt some of the questions were intrusive. Future studies could benefit from the use of a virtual/online platform for the subjective recordings to enhance participant experience. In addition, several cognitive tests could have been excluded on the basis of these pilot data to reduce the time burden of the assessment days.

This study did show direct effects of a physical activity intervention on cognition. Our findings, demonstrating the benefit of physical activity on ToL performance, is supported by the work of Chang et al, 2011^26^ who also found that participants in an exercise group achieved improvements in ToL task scores. These findings indicate that physical activity may have a positive effect on the executive functions of planning and problem solving. Interestingly, a study by Manjunath & Telles, 2001^27^, reported improved performance in the ToL task following yoga training compared with a group who conducted physical exercise. A recent study by Suwabe et al, 2018^28^, showed that acute very light exercise (similar to yoga and tai chi) improved hippocampal memory function and concluded that a modest increase in physical activity in a healthy cohort may improve memory. Our findings showing an improvement in BVMT-R score following the intervention support this work. Further studies are warranted to determine the differential effects of types of physical activity on cognitive function.

Observations in animal models have raised some important questions about the impact of physical activity on neurodegenerative disease processes affecting cognition. In a murine model of Huntington’s disease, light exercise accelerated onset of disease symptoms vs sedentary controls suggesting that exercise is not necessarily beneficial in systems with an impaired nervous system^29^. In addition, in a study evaluating the effects of moderate-high intensive exercise in patients with dementia, it was found that exercise did not slow cognitive impairment^30^. These studies illustrate the importance of cohort selection, the time in which an intervention is implemented, and perhaps how exercise regimens may need to be refined for specific populations.

In our study, on average, % of light and moderate physical activity predicted poorer sleep, as demonstrated by the number of awakenings and sleep efficiency. Our findings contrast with previous data demonstrating positive effects of physical activity on sleep^31,32,33^ and suggests that the improvements we observed on the ToL and BVMT-R tasks were not necessarily mediated through sleep as predicted. It’s important to note that we did not evaluate the time of day that participants conducted most of their physical activity. Various studies have suggested the importance of circadian rhythms and sleep, for example, late night exercise has been linked to sleep difficulties^34^. However, a systematic review conducted by Stutz et al, 2018^35^ concluded that there was no support for evening exercise negatively affecting sleep although sleep-onset latency, total sleep time and sleep efficiency may have been impaired after vigorous exercise ending less than one hour before bedtime. We also cannot ignore the effect of other important biological and psychological factors affected by physical activity such as enhanced vagal modulation, cortisol and growth hormone secretion changes and mood^36^ as well as muscle inflammation/metabolism and parasympathetic activity. A meta-analysis conducted by Kredlow et al, 2015^31^ showed that acute and regular exercise had small beneficial effects on total sleep time and sleep efficiency but no association was found for exercise intensity or aerobic/anaerobic classification. Gabriel & Zierath, 2019^37^ suggested that timing physical activity to coordinate with an individual’s circadian rhythms may be the optimal route for the health benefits. It can be suggested from these results that similar studies in the future would benefit from the inclusion of a lux measurement tool to provide further insight into the potential interaction between time of day and physical activity completion. In addition, further analysis into how we categorise light, moderate and vigorous activity in different populations, and evaluation in a controlled environment would also be a useful avenue for future clinical trials.

## Conclusion

Understanding the relationship between physical activity, sleep and cognition could be an important tool to help combat cognitive impairment in later life. However, the interactions between these factors are not straightforward. In addition to larger sample sizes, individually tailored physical activity programmes, regular feedback to maintain motivation, briefer subjective measures and cognitive assessments (including BVMT-R and ToL tasks) and regular objective outcome measures (using technologies such as actigraphy and lux tools), would be useful to ensure the reliability and relevance of future studies.

### Data Availability

The datasets generated during and/or analysed during the current study are available from the corresponding author on reasonable request.

## Methods

### Ethics Approval

This study was approved by the University of Bristol Research Ethics Committee and conducted in accordance with Good Clinical Practice. All participants provided written informed consent.

### Cohort

Healthy older (≥50) and younger (<50) male and female participants were recruited from the Join Dementia Research website and the ReMemBr group healthy volunteer database. Each participant was pre-screened using the adult pre-exercise screening tool (Exercise and Sports Science *Australia*, Fitness *Australia* or Sports version 11) to evaluate eligibility, including risk factors for physical exercise, sleep disorders and mood disorders. Participants were excluded if a) there was any evidence of significant cognitive, mood or sleep disorders, b) they were taking medications likely to interfere significantly with sleep or cognition, c) they had any significant lifestyle routines that interfered with sleep or cognition i.e. presence of a newborn in the household, d) they were at high-risk for any cardiovascular complications or e) they had any physical problems that would impair mobility.

### Study Design

Participants were enrolled on the study for 14 weeks in total (2 weeks of baseline observations; 12 weeks of intervention). The timeline of the study is illustrated in **Figure 3**. During this time participants made three visits to the Brain Centre at Southmead Hospital in Bristol (UK); at baseline (first contact week 0), midline (end of week 6) and end of visit (end of week 12). Assessments were conducted at each visit as outlined in **Table 4**.

**Figure 3.**
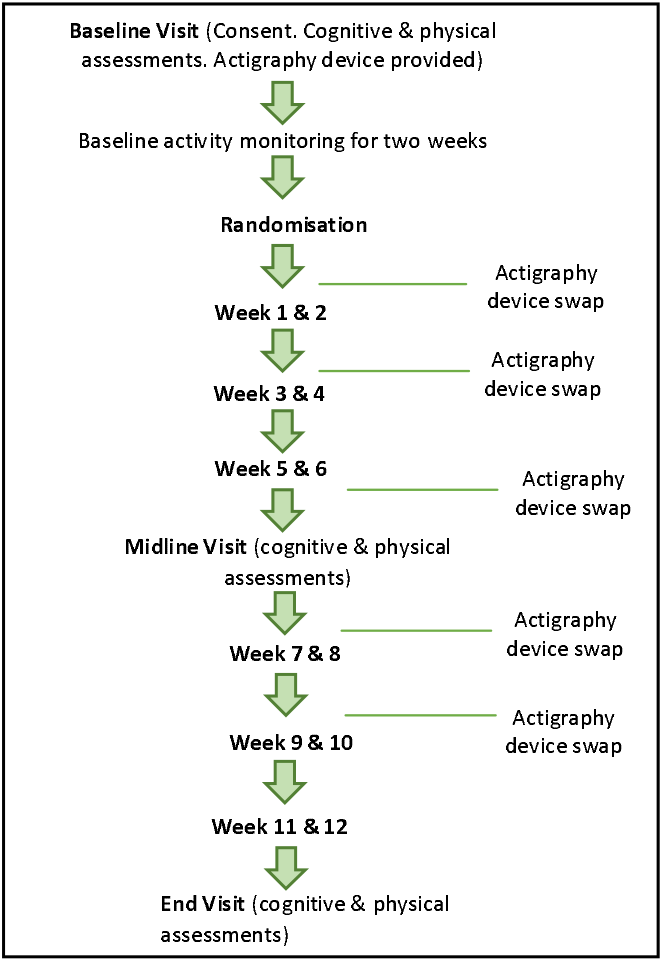
Timeline of events during the study. Participants attended a baseline visit to provide consent and conduct initial physical and cognitive assessments. An actigraph was provided for baseline sleep and activity measurements for the following 2 weeks. At week 3, the participant was randomised to either an exercise or control group and continued to wear the device for 2 week epochs until midline visit. This was repeated until week 12 for the end visit.

**Table 4.** List of assessments for baseline, midpoint and end visits. Participants had physical and cognitive assessments at three points during the study (baseline, midline (end of week 6) and end visit (end of week 12)).

| Physical | Neuropsychology |
| --- | --- |
| Height (baseline visit only) | Years of education |
| Weight (baseline visit only) | DASS |
| Blood pressure (average of three readings) | GDS |
| Pulse oximetry | Profile of Mood States |
| Peak flow reading | Pittsburgh Insomnia Rating scale |
| Resting heart rate | MoCA |
| Chester step test | BVMT-R |
|  | Hopkins Verbal Learning Test-revised (HVLT) |
|  | Serial reaction time (SRT) |
|  | ToL |
|  | Maniken Test of Spatial Orientation and Transformation |

At baseline, all participants were provided with an actigraphy device as well as sleep and activity diaries (Consensus Sleep Diary) to record their routines subjectively. All devices were wrist-worn (n=3, dominant arm; n=20, non-dominant arm). Following baseline observations (end of week 2), participants were randomised to either the intervention or control group using a random number generator. Investigators were blinded to the randomisation. Participants in the intervention group were asked to increase activity by either 4000 steps/day, or by conducting an extra 20-30 min of low-impact physical activity/day, until end of study at week 12. Participants in the control group were asked to not alter their routines from baseline. Due to limited battery life, participants replaced their actigraphy devices every 14 days.

### Actigraph

Sleep and physical activity levels were measured using an actigraphy device provided by ActiGraph Corp (GT9X Link, Actigraph, Pensacola FL), and analysed with ActiLife 6 (ver 6.11.9) software. Compliance was measured from % wear time of the device. Estimates of sleep quality and physical activity parameters were extracted based on the number of counts (accelerometer values used by the actigraphy monitor) for a given epoch. We used 60s epochs, which is consistent with current guidelines for estimating sleep quality^38^. Daily sleep measurements included sleep efficiency (total time in bed/total sleep time x 100), time to first wake after sleep onset (min), number of awakenings and average length of awakening (min). Daily physical activity measurements (data filtered to capture activity between 7am-10pm) included step count, % of time sedentary, % of time spent in light, moderate & vigorous exercise. Cut-off points for sedentary, light, moderate and vigorous activity were used from Neil-Sztramko, 2017^39^ (verified for compatibility with GT9X Link by John et al, 2018^40^, Rowlands et al, 2018^41^ & Montoye et al, 2018^42^). Step count was not included in our analysis due to insufficient evidence of accurate measurement through wrist-worn GTX9 Link monitors. Sleep data were obtained using Sadeh et al, 1994^43^/Cole et al, 1992^44^ algorithms for younger/older participants, respectively.

### Data Analysis

We used the following parameters for sleep and physical activity analysis: sleep efficiency, total sleep time, number of awakenings, % of time spent sedentary, % of time in light physical activity, % of time spent in moderate physical activity and % of time spent in vigorous physical activity. Data were analysed in 2-week epochs. As a representative statistical analysis, we compared baseline with weeks 1 & 2 and weeks 11 & 12 to evaluate initial and sustained effect of intervention, respectively. Multivariate analysis was used with age as covariate and Mauchly’s Test of Sphericity and Huynh-Feldt correction where necessary. As this study is exploratory for feasibility, we quote effect sizes as well as significance levels. Pearson and Spearman’s correlation analysis with Bonferroni correction for multiple comparisons were used as appropriate. GraphPad Prism Version 6.0 (GraphPad Software, La Jolla, CA, USA) and IBM SPSS Statistics for Windows (Version 24.0. Armonk, NY: IBM Corp) software packages were used. Statistical significance was set at p⍰<⍰0.05.

## Contributions

Conceptualization, N.S, E.C; Methodology, N.S, E.C; Investigation, M.S, N.S, A.C, E.G; Formal Analysis, M.S, E.C; Data Curation, Writing - Review & Editing, M.S, E.C; Supervision, E.C Funding Acquisition, E.C.

